# Genomic and transcriptomic analyses of *Heteropoda venatoria* reveal the expansion of P450 family for starvation resistance in spider

**DOI:** 10.1101/2024.07.31.605936

**Authors:** Guoqing Zhang, Yiru Wang, Hongcen Jiang, Yi Wang

## Abstract

**Background:** Spiders generally exhibit robust starvation resistance, with hunting spiders, represented by *Heteropoda venatoria*, being particularly outstanding in this regard. Given the challenges posed by climate change and habitat fragmentation, understanding how spiders adjust their physiology and behavior to adapt to the uncertainty of food resources is crucial for predicting ecosystem responses and adaptability.

**Results:** We sequenced the genome of *H. venatoria* and, through comparative genomic analysis, discovered significant expansions in gene families related to lipid metabolism, such as cytochrome P450 and steroid hormone biosynthesis genes. We also systematically analyzed the gene expression characteristics of *H. venatoria* at different starvation resistance stages and found that the fat body plays a crucial role during starvation in spiders. This study indicates that during the early stages of starvation, *H. venatoria* relies on glucose metabolism to meet its energy demands. In the middle stage, gene expression stabilizes, whereas in the late stage of starvation, pathways for fatty acid metabolism and protein degradation are significantly activated, and autophagy is increased, serving as a survival strategy under extreme starvation. Additionally, analysis of expanded P450 gene families revealed that *H. venatoria* has many duplicated CYP3 clan genes that are highly expressed in the fat body, which may help maintain a low-energy metabolic state, allowing *H. venatoria* to endure longer periods of starvation. We also observed that the motifs of P450 families in *H. venatoria* are less conserved than those in insects, which may be related to the greater polymorphism of spider genomes.

**Conclusions:** This research not only provides important genetic and transcriptomic evidence for understanding the starvation mechanisms of spiders but also offers new insights into the adaptive evolution of arthropods.

## Introduction

Spiders, as widely distributed arthropods, possess remarkable survival abilities and occupy a unique ecological niche in nature [1]. They play dual roles in the food chain as both predators and prey, which underscores their critical importance in ecological systems [2]. For a long time, the abilities of spiders to spin silk and inject venom have been the main features of interest [3–8], particularly their unique ability to produce up to seven distinct types of silk, an unmatched feat in nature [3, 9].

In addition to their predatory tactics involving silk spinning and venom injection, spiders have evolved robust starvation resistance to cope with unstable food supplies. Spiders typically adapt a sedentary lifestyle, waiting for prey while remaining largely motionless, which highlights the significance of the resting metabolic rate throughout their life cycle [10]. Food supply constraints significantly shape the ecology and behavior of spiders, leading to relatively low metabolic rates [11]. The presence of tracheae plays a significant role in spiders, which have well-developed tracheal systems, as most spiders exhibit metabolic rates far below what would be expected on the basis of their body weight, especially those with two pairs of lungs [12]. Other factors, such as sex, lifespan, reproduction, developmental status, type of prey captured, and high anaerobic energy acquisition capabilities, also significantly influence resting and active metabolic rates. For example, spiders with a lifespan exceeding one year have lower metabolic rates than those with a one-year life cycle [12, 13]. Cellular autophagy is equally crucial for maintaining energy homeostasis during periods of starvation [14]. Research has shown that the evolution of social spiders is linked to nutritional metabolism and autophagy, which regulate metabolic processes and mitigate the threat of cannibalism to ensure an adequate energy supply [15].

Starvation resistance is an adaptive trait evolved by organisms to survive in environments with food scarcity. The starvation resistance of spiders allows them to extend their survival time under conditions of prey scarcity through various physiological mechanisms, such as reducing metabolic rates, decreasing activity levels, and utilizing stored energy. Additionally, the starvation resistance of spiders may be associated with specific behavioral adaptations, such as alterations in predation strategies, optimization of energy allocation, and improvement of the timing of reproductive investment [16]. The starvation resistance of spiders, as a core component of their survival strategy, not only directly impacts individual survival rates but also profoundly influences energy flow and material cycling within ecosystems [16]. As global climate change accelerates and habitat fragmentation intensifies, understanding how spiders adapt their physiological and behavioral strategies to cope with the unpredictability of food resources is crucial for predicting ecosystem responses and adaptability.

Research on plant stress resistance is plentiful and has focused primarily on drought resistance, salt tolerance, chilling tolerance, and other biotic stresses [17–21]. In contrast, studies on animal stress resistance are relatively rare, and the starvation resistance of spiders is highly important in research on ecological adaptation. With the continuous advancement of biological research methods and the development of sequencing technologies [22, 23], we have the opportunity to investigate the mechanisms of spider starvation resistance from molecular, physiological, and behavioral ecological perspectives, as well as the ecological and evolutionary significance of this phenomenon. *H. venatoria* is a huntsman spider and is characterized by their well-developed limbs and extremely fast movement. *H. venatoria* does not spin webs; it is known for hunting live insects with exceptional agility and speed during the night. However, these findings indicate that *H. venatoria* does not feed in a stable manner and often experiences periods of starvation. Additionally, the average lifespan of male *H. venatoria* is 465 days, whereas females live for approximately 580 days [24], both clearly surpassing one year; these spiders can therefore be classified as spiders with a low metabolic rate [12]. To date, research on *H. venatoria* has focused mainly on its venom [25–28], and there is no high-quality genome sequence available for further exploration of the molecular mechanisms underlying its starvation resistance. Therefore, this study aimed to investigate the expression of functional genes associated with starvation resistance in *H. venatoria* through genomic and transcriptomic data, explore its response strategies to environmental changes, and outline future research directions, with the goal of providing a scientific basis for the conservation of biodiversity and the maintenance of ecosystem functions.

## Results

### High-quality genome assembly and annotation of *H. venatoria*

The female *H. venatoria* spider has 22 chromosomes in its haploid set (2n=44) [29]. To assess the complexity of the *H. venatoria* genome, we initially sequenced the female *H. venatoria* genome via Illumina technology and obtained approximately 184 Gb of raw data. K-mer analysis revealed that the genome size was 5.36 Gb, with a repeat proportion of 46.3% and a heterozygosity rate of 0.96% (Fig. S1A). These findings suggest that *H. venatoria* possesses a large genome with high repeat content and heterozygosity, presenting challenges in genome assembly.

To achieve high-quality genome assembly for *H. venatoria*, we employed HiFi sequencing, which resulted in approximately 127 Gb of raw sequencing data. The initial assembly yielded a 5.95 Gb contig genome, with an N50 of 2.4 Mb. Using Hi-C for contig mounting, we ultimately obtained a genome consisting of 22 chromosomes, with a scaffold N50 of 253.94 Mb and a genome size of 5.52 Gb (Table 1), which was closely aligned with the 5.37 Gb obtained by genome survey. The Hi-C map clearly demonstrated high continuity in the chromosome assembly (Fig. 1A). BUSCO analysis revealed an assembly completeness of 96.3%, with only 4.0% duplicated BUSCOs (Table 2), indicating that the high-quality genome assembly was suitable for subsequent analyses.

**Figure 1:**
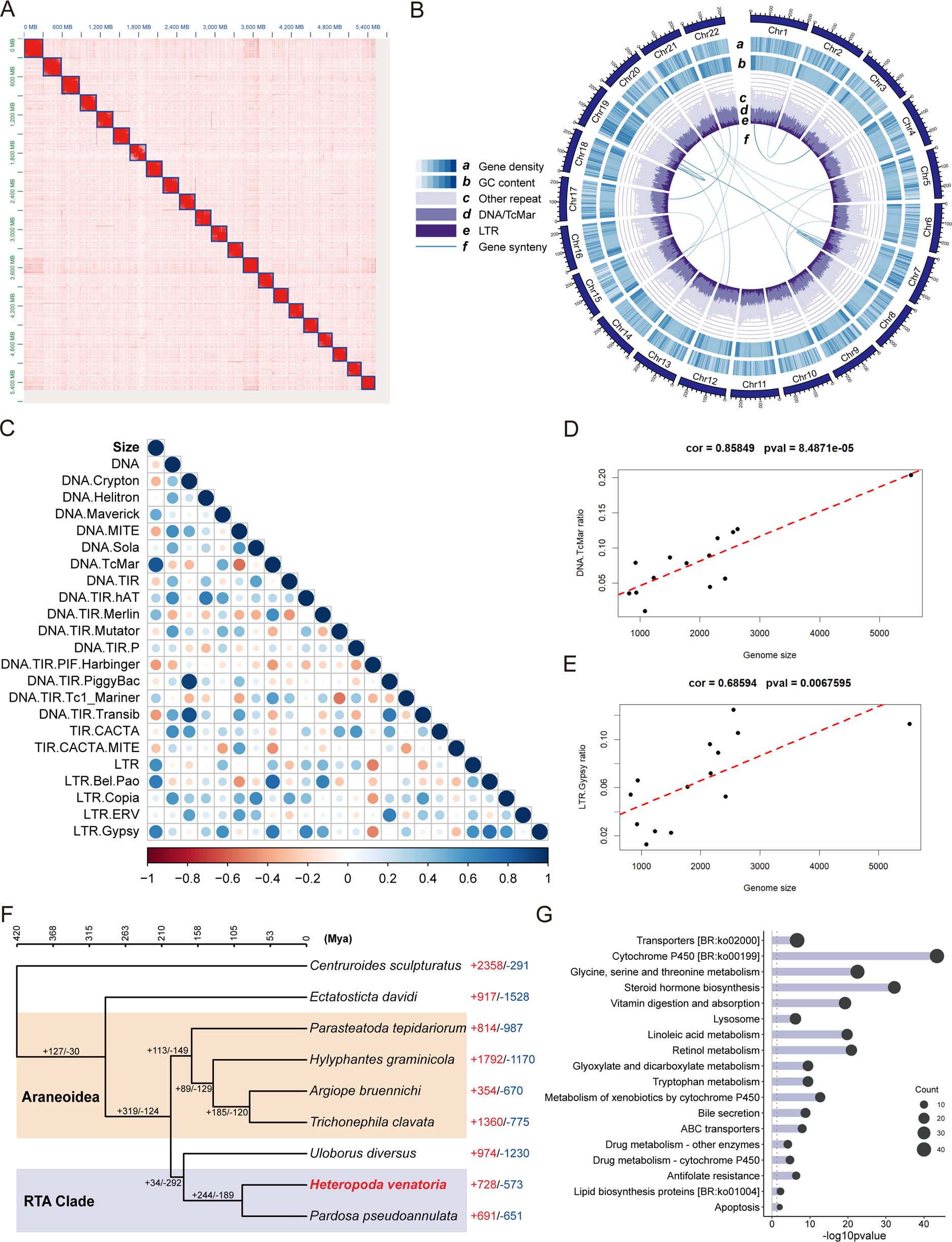
Chromosomal-scale genome assembly and genomic characteristics of *Heteropoda venatoria*. (A) Hi-C assembly map of *H. venatoria*. (B) Circular diagram depicting the genomic features of *H. venatoria*. (C) Correlations between genome size and the prevalence of different types of repeats. (D, E) Linear relationships of DNA.TcMar and LTR.Gypsy with genome size. (F) Phylogenetic tree of a scorpion and seven spider species, along with the contraction and expansion of gene families. (G) KEGG functional enrichment of the expanded gene families in *H. venatoria*.

**Table 1:**
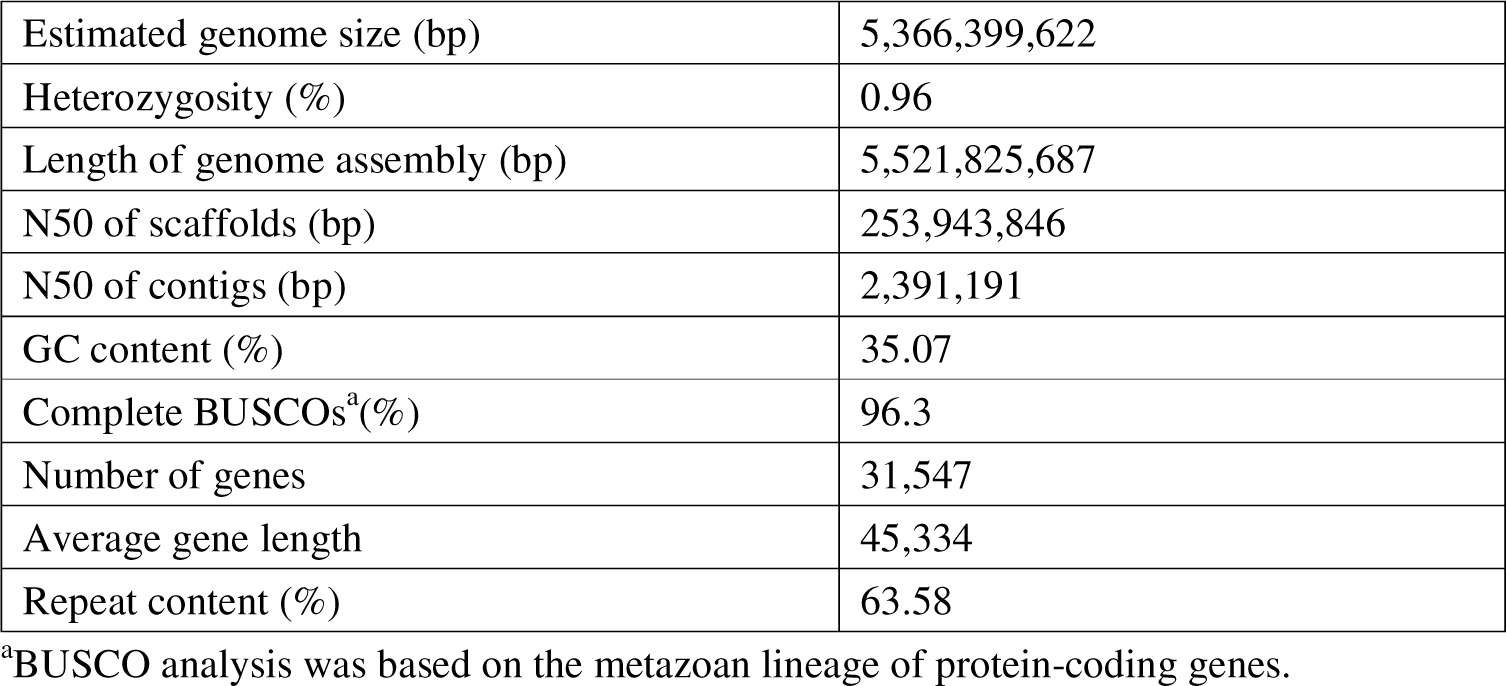
Characteristics of the *Heteropoda venatoria* genome assembly.

|  |  |
| --- | --- |
| Estimated genome size (bp) | 5,366,399,622 |
| Heterozygosity (%) | 0.96 |
| Length of genome assembly (bp) | 5,521,825,687 |
| N50 of scaffolds (bp) | 253,943,846 |
| N50 of contigs (bp) | 2,391,191 |
| GC content (%) | 35.07 |
| Complete BUSCOs<sup>a</sup>(%) | 96.3 |
| Number of genes | 31,547 |
| Average gene length | 45,334 |
| Repeat content (%) | 63.58 |

<sup>a</sup>BUSCO analysis was based on the metazoan lineage of protein-coding genes.

**Table 2:** BUSCO analysis of the *Heteropoda venatoria* genome assembly and annotation.

|  | Genome assembly | Genome annotation |
| --- | --- | --- |
| Complete BUSCOs<sup>a</sup>(%) | 96.30% | 94.30% |
| Complete and single-copy BUSCOs | 92.30% | 90.60% |
| Complete and duplicated BUSCOs | 4.00% | 3.70% |
| Fragmented BUSCOs | 1.70% | 1.50% |
| Missing BUSCOs | 2.00% | 4.20% |

<sup>a</sup>BUSCO analysis was based on the metazoan lineage of protein-coding genes.

In terms of genome annotation, we identified 31,547 genes with an average length of 45 kb (Table 1), achieving a completeness rate of 94.3%. The functional annotations included GO terms for 13,857 genes and KEGG pathways for 9,488 genes. With respect to repeat annotation, repeat sequences accounted for 63.58% of the genome (Table 1), which is significantly greater than the proportion of repeats estimated by K-mer analysis (Supplementary Fig. S1A). We hypothesize that this discrepancy might be due to the shorter read lengths from Illumina sequencing, which could lead to underestimation of the levels of tandem repeats or long interspersed elements. The impact of repeat sequences on genome size is significant [30–34]. Given the relatively high genome size and proportion of repeat sequences in *H. venatoria* compared to other spiders, we additionally collected genomic data from 13 other spider species and identified their repeat sequences. Analysis of repeat sequences across 14 spider genomes revealed a strong correlation between genome size and the presence of TcMar and LTR elements (Fig. 1C-E). Notably, these two types of repeat sequences also had the highest prevalence in *H. venatoria*. We speculate that during the evolution of spider genomes, two types of repeat sequences, TcMar and LTR sequences, had a significant impact on the size of spider genomes. Interestingly, we found that in *H. venatoria* chromosomes, regions with a high proportion of repeats also presented an increase in GC content (Fig. 1B). We speculate that these repeat sequences, which are rich in GC, are subjected to selective pressure due to their structural stability or stronger binding to specific proteins, leading to an increase in the GC content in the repeat sequence regions.

### Gene family expansion and contraction

We gathered genomic data and annotations for one scorpion and seven chromosome-level spider genomes using the scorpion as an outgroup [35–42]. Using the maximum likelihood method, we constructed a phylogenetic tree encompassing these eight arachnid species. Phylogenetic analysis revealed that *H. venatoria* is a member of the RTA clade, which is consistent with recent research findings [43]. Additionally, through the application of CAFE5 for the analysis of gene family expansion and contraction, we found that *H. venatoria* has expanded to a total of 728 gene families (Fig. 1F and Supplementary Table S4). To elucidate the functional implications of these expanded families, we performed functional enrichment analysis on the genes associated with these families that had undergone significant expansion. Our findings revealed that pathways related to lipid metabolism, including cytochrome P450 [BR:ko00199], steroid hormone biosynthesis, linoleic acid metabolism, and lipid biosynthesis proteins [BR:ko01004], were significantly enriched in *H. venatoria* (Fig. 1G). Spiders generally exhibit strong starvation resistance, and *H. venatoria*, a hunting spider, relies on its well-developed legs for hunting rather than on webs, making its feeding more unpredictable. Therefore, we preliminarily speculate that *H. venatoria* frequently experiences starvation, and this formidable starvation tolerance may be associated with the expansion of gene families related to lipid metabolism pathways within its genome.

### Transcriptome design for starvation resistance in *H. venatoria*

To further investigate the reasons behind the exceptional starvation resistance of *H. venatoria*, the samples of *H. venatoria* were subjected to starvation treatments were divided into six groups according to duration of treatment: CK (measured after 3 hours of feeding), 2 W (2 weeks after feeding), 4 W (4 weeks after feeding), 8 W (8 weeks after feeding), 14 W (14 weeks after feeding), and 19 W (19 weeks after feeding). Transcriptome sequencing was performed on both the whole-body and fat body tissues of the spiders, resulting in 12 sets of samples (each with three replicates). PCA revealed that the expression profiles of the fat body transcriptome were more closely correlated with the duration of starvation in *H. venatoria* (Fig. 2A), whereas the whole-body transcriptome showed some overlap between different treatments (Supplementary Fig. S1A), likely due to the inclusion of numerous tissues, which resulted in tissue-specific variations overshadowing treatment effects. After removing outlier samples from the whole-body transcriptome, the results revealed that, except the 14 W and 19 W samples, which showed obvious differences from the other samples, the remaining samples presented relatively minor expression variations (Fig. 2B).

**Figure 2:**
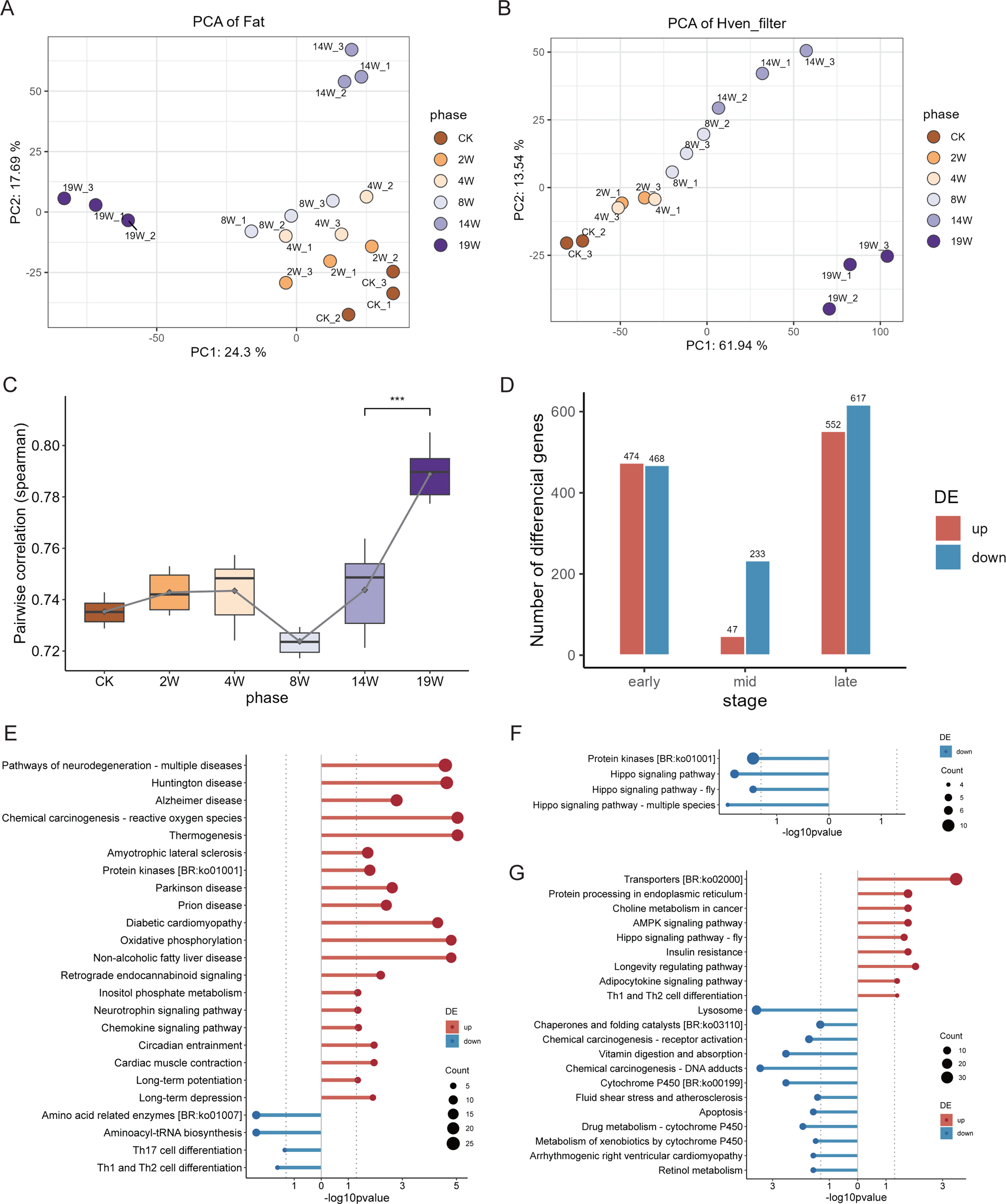
Transcriptomic analysis results of the fat body and whole-body responses to starvation resistance in *Heteropoda venatoria*. (A, B) Principal component analysis (PCA) results of the starvation resistance-related transcriptome. (C) Correlation analysis of the transcriptomes of the fat body and whole-body across various stages of starvation resistance. (D) Number of DEGs in the fat body transcriptome during the early, middle, and late stages of starvation resistance. (E, F, G) KEGG functional enrichment results for DEGs in the fat body transcriptome during the early, middle, and late stages of starvation resistance.

According to the PCA of the fat body expression, the samples were clustered into four distinct groups (Fig. 2A). Notably, the 4 W and 8 W samples clustered closely, with CK and 2 W being relatively proximal, while 14 W and 19 W were markedly divergent. We hypothesize that 14 W and 19 W represent the later stages of starvation, when *H. venatoria*’s overall gene activity is heightened, leading to greater transcriptomic differences in these samples. For subsequent analysis, we divided the starvation process into three phases: early starvation (CK and 2 W), middle starvation (4 W and 8 W), and late starvation (14 W and 19 W). Both the fat body and whole-body expression analyses revealed that the 19 W samples were obviously different from the other samples (Fig. 2A, B), suggesting that at 19 weeks, *H. venatoria* reached the limit of its starvation period, with distinct functional expression occurring within its body. Therefore, in addition to the three main phases, the 19 W samples were analyzed separately.

### Differential transcriptomic analysis during early, middle, and late stages of starvation in *H. venatoria*

In our study of the fat body transcriptome of *H. venatoria* during three distinct stages of starvation (early, middle, and late), we observed the following expressing patterns:

During the early stage of starvation (from CK to 2 W), many genes, specifically those involved in oxidative phosphorylation and thermogenesis pathways, were upregulated (Fig. 2E). These findings indicate that during the early starvation stage, energy metabolism in *H. venatoria* occurs regularly, with sufficient supply of energy. In the middle starvation phase (from 4 W to 8 W), the number of DEGs was the lowest. Some of the downregulated genes were significantly enriched in the Hippo signaling pathway, a pathway closely associated with sugar and lipid metabolism (Fig. 2F). In liver cells, the core kinase LATS2 of the Hippo signaling pathway binds with and inhibits the activity of SREBP, a key enzyme in cholesterol synthesis. Liver-specific deletion of Lats2 results in overactivation of SREBP, leading to abnormal lipid accumulation in the liver [44]. However, other studies have shown that in brown adipose tissue, YAP/TAZ can promote thermogenesis and reduce fat accumulation in the body by upregulating the expression of UCP1. Knocking out YAP/TAZ in brown adipose tissue resulted in increased body weight and body fat percentage in mice, as well as impaired glucose tolerance and insulin sensitivity [45]. These findings indicate that the Hippo signaling pathway involves tissue-specific regulation of lipid metabolism. We hypothesize that during the middle starvation phase, *H. venatoria* may reduce its cellular glucose uptake and utilization to adjust to the food-scarce environment. During the late starvation phase (from 14 W to 19 W), pathways involved in protein transport and processing within the endoplasmic reticulum become particularly active, potentially owing to the insufficient supply of glucose-based energy and a primary reliance on fat for energy. This is also indicated by the activation of the AMPK signaling pathway, insulin resistance, and the adipocytokine signaling pathway. Additionally, pathways related to cell proliferation and autophagy, such as the lifespan regulation pathway, choline metabolism in cancer, the Hippo signaling pathway in Drosophila, and Th1 and Th2 cell differentiation, become more active (Fig. 2G). We speculate that due to severe energy deficiency, the late starvation phase could be a period of increased cellular autophagy. Due to the reduced number of nonessential cells, energy expenditure decreases, and some energy is derived from the breakdown of cells.

As PCA revealed a strong correlation between the expression profiles of the adipose tissue transcriptome and starvation tolerance duration in *H. venatoria*, we conducted a weighted gene co-expression network analysis (WGCNA) of the fat body transcriptome [46]. Clustering of the 18 fat body samples via WGCNA yielded a total of nine modules, including the grey module (Supplementary Fig. S3A). Notably, the blue and brown modules exhibited significant correlations with the entire starvation process (Supplementary Figs. 4C, 4D). These two modules are hypothesized to play a dominant role in starvation tolerance. In the blue module, the majority of genes presented increased expression with prolonged starvation duration (Supplementary Fig. S4A), whereas a subset of genes were downregulated. Conversely, most genes in the brown module presented the opposite trend (Supplementary Fig. S3B). Functional enrichment analysis of the genes in these two modules revealed that, in addition to the previously mentioned AMPK signaling pathway, insulin resistance and adipocytokine signaling pathway, which are active in the later stages, the citrate cycle and PPAR signaling pathway also exhibited heightened activity during the late stages of starvation. We speculate that the upregulation of the citrate cycle is aimed primarily at reducing glucose dependency and increasing fatty acid oxidation while also modulating the metabolism of certain amino acids. The elevated expression of PPARα and PPARδ in the PPAR signaling pathway promotes fatty acid transport and oxidation, thereby regulating energy balance.

### Final starvation stage in *H. venatoria*

The PCA results from both the fat body and whole-body transcriptomes indicated that *H. venatoria* transcriptome at 19 weeks of starvation was markedly distinct from that at other stages (Fig. 2A, B). Consequently, we conducted a differential analysis of the transcriptome at 19 weeks. Differential analysis of fat body tissue at 19 weeks identified 612 upregulated genes and 647 downregulated genes (Supplementary Fig. S7A and Supplementary Table S5). The functional enrichment results revealed that only transporters and autophagy were significantly upregulated at 19 weeks, whereas energy-consuming pathways such as DNA replication and cell cycle were essentially inactive (Supplementary Fig. S7B).

WGCNA of the fat body revealed that the lysosomal pathway is enriched in multiple modules in *H. venatoria*, indicating that the expression of different functional lysosomes during starvation in *H. venatoria* is also distinct. Specifically, the six genes in the blue module and the eleven genes in the brown module exhibited similar expression patterns during starvation, essentially showing continuous downregulation (Supplementary Fig. S4A-D), and the functional annotation results for these 17 genes indicated that they mainly encoded glycosidases, lipases, and proteases in the lysosome (Table S3). These findings suggest that during starvation in *H. venatoria*, these enzymes perform continuous proteolysis and recycle some cellular components, providing a steady energy supply for *H. venatoria*. However, as the number of cellular components composed of sugars, lipids, and proteins decreases, this regulatory process also gradually weakens until it becomes almost completely silent at 19 weeks.

Interestingly, 24 genes of the lysosomal pathway in the turquoise module exhibited a sharp increase in expression at 14 weeks (Supplementary Fig. S4E, F and Supplementary Table S2). Compared with the aforementioned 17 genes, this group included more proteases and, more importantly, it included three genes related to sulfatases (Table S3). The primary function of lysosomal sulfatases is to degrade sulfated glycosaminoglycans and glycolipids. This finding indicates that in the final stages of starvation, *H. venatoria* gradually breaks down its internal connective tissues (sinusoidal tissues) in skeletal and joint areas to provide energy for its last phase of life.

In contrast, the whole-body transcriptome at 19 weeks showed a substantial increase in upregulated genes, which was significantly greater than that in any other period (Fig. 3A). The functional enrichment results revealed that, in addition to the upregulation of pathways such as transporters and lysosome pathways, pathways such as cytochrome P450 [BR:ko00199], steroid hormone biosynthesis, and linoleic acid metabolism pathways also exhibited significant upregulation (Fig. 3B). These pathways have undergone notable gene family expansion in *H. venatoria* and are all related to lipid metabolism.

**Figure 3:**
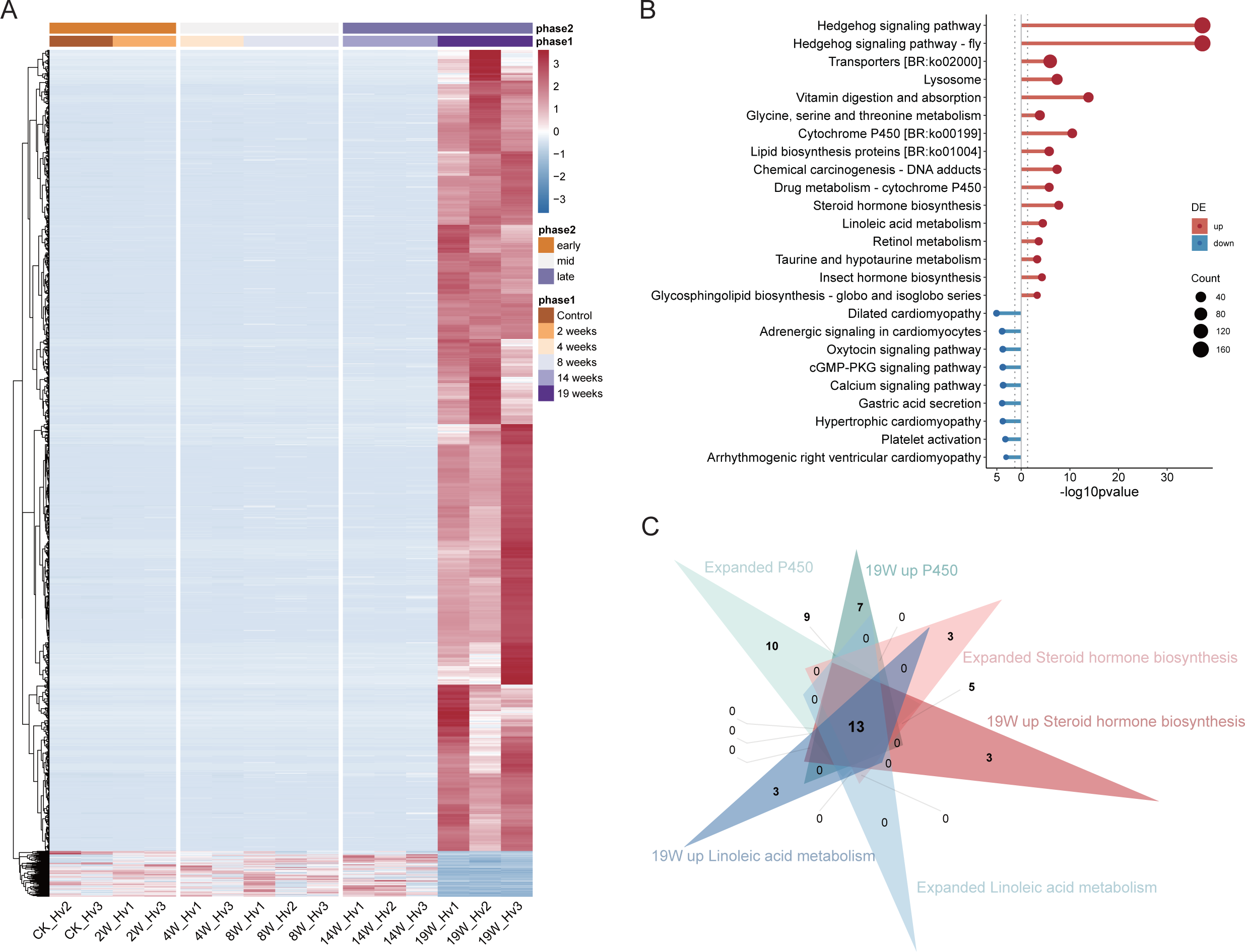
Whole-body transcriptome analysis of *Heteropoda venatoria* during a 19 W starvation period. (A, B) Heatmap and KEGG pathway enrichment analysis of DEGs in the whole-body transcriptome of *H. venatoria* at 19 W starvation period. (C) The overlap of genes enriched in the cytochrome P450, steroid hormone biosynthesis, and linoleic acid metabolism pathways within the expanded gene families of *H. venatoria* and the same pathways identified at 19 W among upregulated genes of the whole-body transcriptome related to starvation resistance.

An overlap analysis of the genes enriched in these three expanded pathways and the genes upregulated at 19 weeks revealed 13 shared genes (Fig. 3C). Functional annotation of these 13 genes revealed that they are all associated with P450 (cytochrome P450), and interestingly, all are located on Chr4. We hypothesize that P450 genes play crucial roles in the starvation response of *H. venatoria*. Consequently, we conducted an identification analysis of P450 genes in *H. venatoria* and seven other spider species.

### P450 genes in eight spiders

The identification of P450 genes across eight spider species revealed a total of 714 P450 genes encoding 848 P450 proteins. Phylogenetic tree construction revealed that spider P450 genes can be classified into the CYP2 clan, CYP3 clan, CYP4 clan, and mitochondrial clan (Fig. 4A and Supplementary Figure S8), which is consistent with findings in most arthropods [47]. Notably, all 13 genes from the three enriched pathways mentioned earlier belong to the CYP3 clan. A phylogenetic tree constructed for the CYP3 clan of the eight spider species showed that the 23 CYP3 clan genes from Chr4 of *H. venatoria* (representing 25 proteins) all clustered on the same branch. Additionally, two other genes from Chr4 of *H. venatoria* and the CYP3 genes from five other spider species were grouped together on a separate branch (Fig. 4B). Interestingly, these 25 genes are located within a region less than 2.5 Mb in size, leading us to preliminarily deduce that the branch on which these genes reside likely represents a single subfamily. This finding also indicates considerable expansion of the CYP3 subfamily in *H. venatoria*. We therefore conducted a synteny analysis of the proteins on this branch and the protein Ectatosticta_davidi_00009731_1 from *E. davidi*, which is the closest relative to this branch.

**Figure 4:**
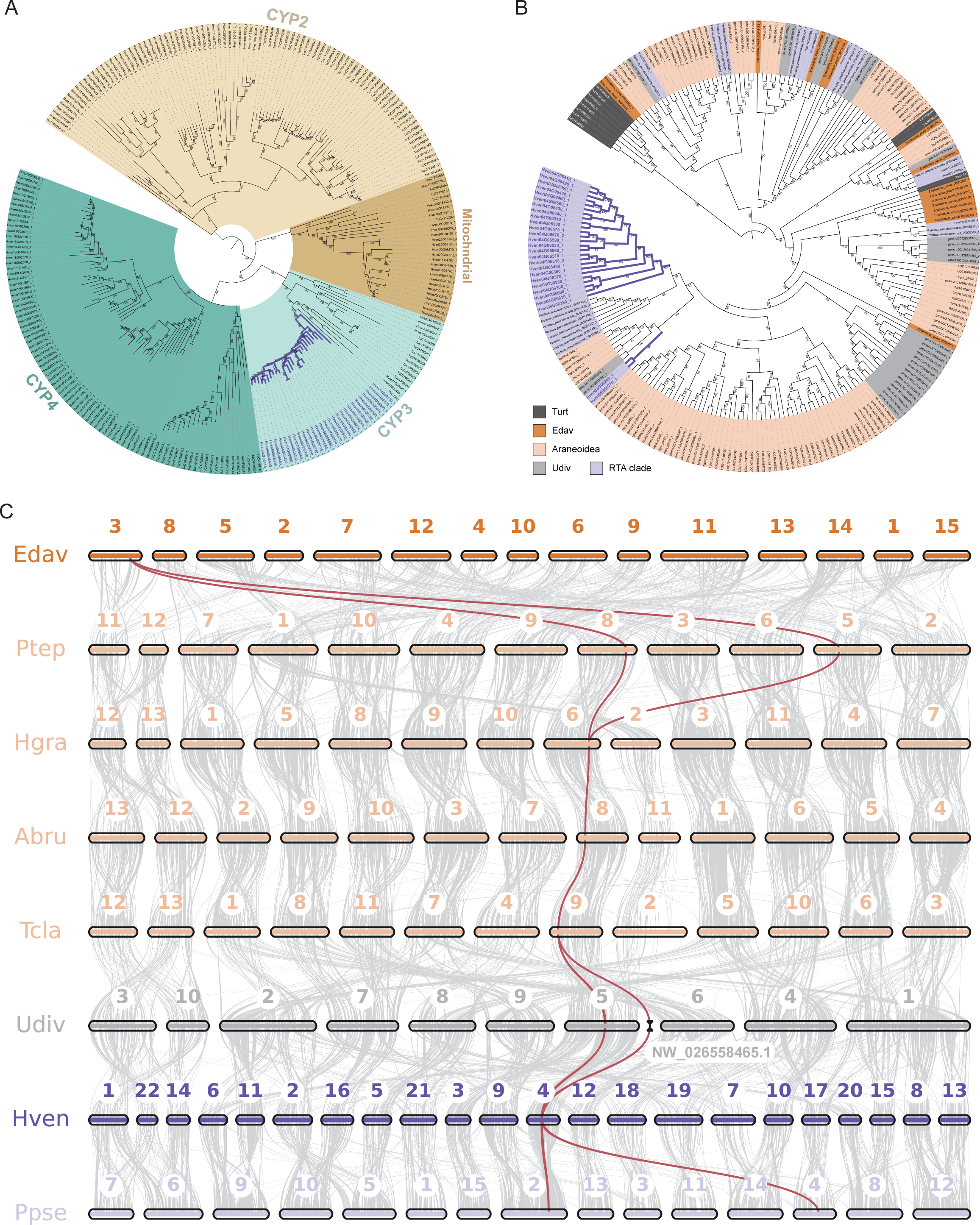
Evolutionary analysis of cytochrome P450 genes. (A) Phylogenetic tree of P450s in *Heteropoda venatoria* and *Tetranychus urticae*. (B) Phylogenetic tree of the CYP3 clan genes in eight spider species and *T. urticae*. (C) Collinearity relationships of a subset of CYP3 clan genes in eight spider species.

Synteny analysis revealed that *H. venatoria* has an increased number of chromosomes due to extensive chromosomal fragmentation. In *Uloborus diversus*, members of this subfamily are located on Chr5 and an unanchored scaffold. After chromosomal fragmentation, the main fragments of *U. diversus* Chr5 corresponded to Chr3 and Chr4 in *H. venatoria*, wherein significant gene duplication occurred on Chr4, leading to the expansion of the CYP3 subfamily (Fig. 4C).

Research has shown that a subfamily within CYP3, specifically CYP3A, can inhibit the metabolic rate of glucose in female mice, leading to an increase in fat [48]. As most genes of the CYP3 subfamily in *H. venatoria* are expressed in the fat body during various stages of starvation, we speculate that the numerous copies of the CYP3 subfamily genes maintain relatively low energy metabolism. This adaptation may allow *H. venatoria* to survive for extended periods without feeding. To further investigate the CYP3 subfamily in *H. venatoria*, we conducted additional sequence analyses.

### Conserved domains in P450s

Insect P450s are known to contain five conserved motifs: the helix C motif (WxxxR), the helix I motif (GxE/DTT/S), the helix K motif (ExLR), the PERF motif (PxxFxPE/DRE), and the heme-binding motif (PFxxGxRxCxG/A) [47, 49].

In *H. venatoria*, the WxxxR and PFxxGxRxCxG/A motifs are highly consistent with those in insects (Fig. 5A). However, the ExxR motif in spiders differs from the insect ExLR motif in that the third position of the motif was less conserved, although it still predominantly features L, with significant occurrences of Q and other amino acids (Supplementary Fig. S11 and Fig. S12). The helix I motif (GxE/DTT/S) of insects was also less conserved at the third and fourth positions in *H. venatoria*, leading to the renaming of this motif as GxxTx. Similarly, the PERF motif (PxxFxPE/DRE) of insects was also less conserved at the first, fourth, seventh, and ninth positions in *H. venatoria*, resulting in the renaming of this motif as P/AxxF/YxPxRF/W (Fig. 5A and Supplementary Fig. S11). Compared with the conserved motifs in insects, the motifs in spiders exhibit greater variability at many positions. Additionally, the syntenic relationships among spider genomes reveal extensive chromosomal breakage and fusion events (Fig. 4B), which undoubtedly increase the number of genomic polymorphisms in spiders. This likely contributes to the reduced number of conserved sites in spider P450 genes.

**Figure 5:**
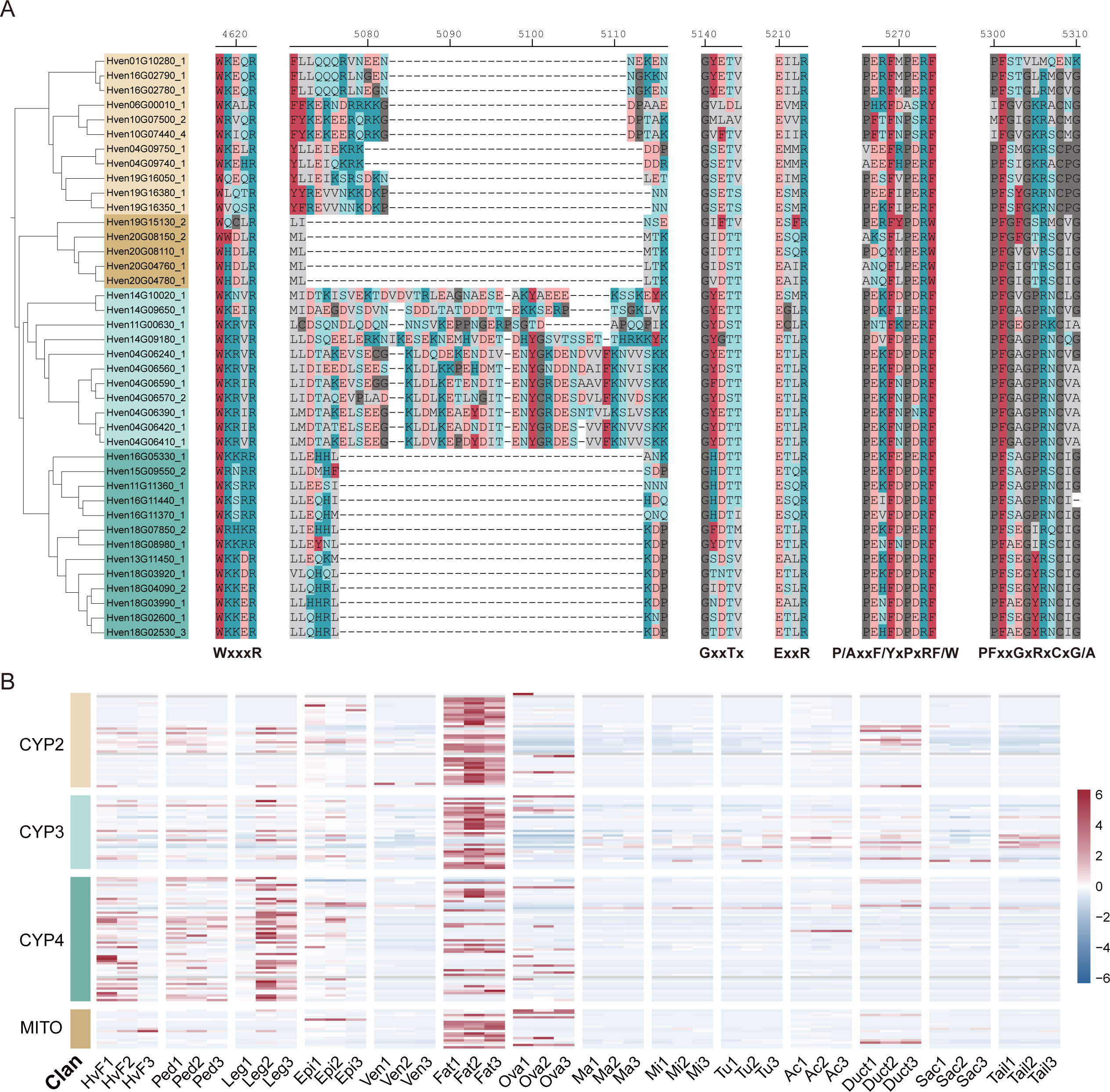
Analysis of conserved motifs in P450 genes and their expression across various tissues in *Heteropoda venatoria*. (A) The five common conserved motifs of the partial P450 genes and a specific sequence of CYP3 genes in *H. venatoria*. (B) P450 genes expression of fat body and other tissues in *H. venatoria*.

### High expression of P450 genes in the fat body

To further investigate the function of P450 genes, we analyzed the expression profiles of four P450 clans in various tissues of *H. venatoria*. The heatmap indicates that the CYP2, CYP3, and mitochondrial genes are predominantly expressed in the fat body, while the CYP4 genes are expressed not only in the fat body but also at significant levels in the pedipalps and legs (Fig. 5B). This finding suggested that, in addition to the CYP3 genes, the CYP2 and mitochondrial genes also play crucial roles in energy metabolism. For comparison, in *Pardosa pseudoannulata*, which belongs to the same RTA branch, the CYP2 and CYP3 genes are also highly expressed in the fat body, but the mitochondrial genes are not significantly expressed in the fat body [50]. Since transcriptomic data for the fat body of *Trichonephila clavata* are available, we also examined the expression of P450 genes in the fat body and other tissues of *T. clavata*. The analysis revealed that the CYP2, CYP3, and some CYP4 genes are highly expressed in the fat body in this species, whereas mitochondrial genes are predominantly expressed in the ovary (Supplementary Fig. S10). These findings indicate that the majority of P450 genes in spiders indeed function in the fat body, allowing spiders to maintain a low-energy metabolic state and increasing their starvation resistance and thus survival in complex predatory environments.

## Discussion

In summary, our study is the first to systematically analyze gene expression differences in *H. venatoria* during various stages of starvation resistance, revealing metabolic pathways and signaling pathways associated with starvation tolerance. Through comparative analysis of the whole-body transcriptome and fat body transcriptome, we found that changes in the fat body transcriptome correlated strongly with starvation duration, suggesting that the fat body may play a crucial regulatory role in the starvation response of *H. venatoria*. Spiders have a large abdomen containing abundant fat bodies, indicating that they have ample energy reserves. Analysis of the fat body transcriptome of *H. venatoria* also showed that the fat body plays a significant role in energy metabolism throughout starvation, participating in the initial strong glucose metabolism, in reduced glucose uptake and utilization by cells in the middle stage, and in increased AMPK signaling and autophagy in the later stage.

Although the fat body of *H. venatoria* provides ample energy reserves, allowing a starvation period of nearly five months, it must maintain a relatively low metabolic rate to slow energy consumption. Interestingly, we found that some P450 families are expanded in *H. venatoria*, and most P450 genes are more highly expressed in the fat body than in other tissues (Fig. 5B). During the starvation experiment, most P450 genes were expressed at various stages in the fat body (Supplementary Fig. S10A). However, the results from whole-body transcriptome analysis were markedly different, with P450 genes showing higher expression at 19 weeks than at other times (Supplementary Fig. S10B). On the basis of existing research showing that some P450 families can reduce metabolic rates or participate in the regulation of lipid metabolism [48, 51–53], we hypothesize that during most starvation periods in *H. venatoria*, P450 genes are expressed primarily in the fat body to inhibit metabolic rates. As starvation progresses to the final stage (19 W), when energy reserves in the fat body are depleted, various *H. venatoria* tissues rely primarily on autophagy to function, with P450 gene expression triggered in most tissues to suppress metabolic rates. In addition to being highly expressed in the fat body in *H. venatoria*, P450 genes are also predominantly expressed in fat bodies in *P. pseudoannulata* and *T. clavata*. These findings indicate that the role of P450 genes in the inhibition of metabolic rates is a common phenomenon in spiders, not only in *H. venatoria*. Owing to the well-developed P450 families in spiders and to their dormancy habit [10], spiders have a low metabolic rate, allowing them to endure long periods of starvation, with the expansion of P450 families in *H. venatoria* resulting in starvation resistance.

The phylogenetic tree of spider P450 genes indicates that many spiders, after acquiring P450 genes from their ancestors, have generated numerous copies within their genomes to enhance their functions. For example, the 23 CYP3 clan genes on *H. venatoria* Chr4 (encoding 25 proteins) all cluster on the same branch, and synteny analysis results showed that they originated from the *U. diversus* CYP3 clan genes. However, how these CYP3 clan genes in *H. venatoria* were duplicated and whether this duplication is related to complex chromosomal breakage and fusion phenomena in spiders require further research and exploration.

To increase the efficiency of fatty acid and amino acid transport, amino acid-related enzymes and transporters are upregulated synchronously during starvation in *H. venatoria*, promoting the transport of fatty acids and amino acids for energy. This may constitute an optimized energy utilization strategy, allowing the body to preserve crucial protein synthesis mechanisms in extreme environments. The gradual upregulation of the insulin-resistant TCA cycle indicates that fats and proteins become the primary energy sources in the middle to late stages of starvation. This metabolic reorganization reveals adaptive regulation in *H. venatoria* under extreme energy constraints.

Finally, autophagy plays a crucial role in the cell response to starvation and other stress conditions. In *H. venatoria*, the lysosomal pathway is enriched in multiple modules, indicating variability in expression regulation during starvation. This process weakens as starvation persists, suggesting a decreasing supply of substances available for degradation within cells. Moreover, the upregulation of genes involved in sulfatase activity in the lysosomal pathway indicates that *H. venatoria* may begin to break down its internal connective tissues for energy under extreme starvation. Understanding this physiological regulatory mechanism under severe energy constraints may be important for understanding the survival strategies and adaptive limits of arthropods.

## Methods

### Genome and Hi-C sequencing

We selected mature female *Heteropoda venatoria* whole tissue for library construction. Using PacBio HiFi sequencing for library preparation, we obtained high-quality full-length DNA for the entire genome. The DNA was fragmented using Megaruptor and then sorted by Sage ELF for 13-16K fragments, followed by adapter ligation to obtain a SMRTbell library [22].

Next, we prepared the Hi-C library [54]. First, formaldehyde was used to fix DNA-protein or protein-protein complexes that were naturally cross-linked or spatially close within *H. venatoria* cells. Chromatin was subsequently digested and separated using the restriction enzyme DpnII, with end repair and biotin labeling of the fragment ends. DNA ligase was used to connect the ends, forming a circular chimeric molecule. These circular molecules were purified and then fragmented into DNA fragments. Finally, the biotin-labeled target DNA fragments were captured using a biotin precipitation technique, and DNA fragments of appropriate size were selected to establish the Hi-C library, which was then sequenced using the DNBseq platform for paired-end sequencing.

### Transcriptome sample processing

We selected mature female adult *H. venatoria* for this study. Preliminary tests of the starvation tolerance cycle revealed that the starvation period of *H. venatoria* was approximately 18 to 20 weeks. Therefore, the samples were divided into six groups based on starvation stage: just after feeding (CK), 2 weeks after feeding (2 W), 4 weeks after feeding (4 W), 8 weeks after feeding (8 W), 14 weeks after feeding (14 W), and 18-20 weeks after feeding (determined by the spider’s condition). The environmental temperature was set at 22±2°C during the day (9:00-19:00) and 16±2°C at night (19:00-next day at 9:00), with a humidity of 70±10% and a natural light cycle. Both the adipose tissue and the entire spider were sampled from each group for transcriptome sequencing. Before the formal starvation experiment, preliminary processing was necessary to ensure that there were enough samples that could feed within the same time frame. After the samples were obtained, unrestricted feeding was allowed during the first week, and feeding was stopped in the second week for one week to ensure that most of the spiders were in a state of hunger. On the first day of the third week, formal feeding commenced, and samples were selected for subsequent experiments within three hours. The spiders were divided into six groups, with eight spiders in each group (including two as backups). In the first group (CK), the adipose tissue and the entire spider were sampled from three samples each three hours after feeding; in the second group, the tissues were sampled after two weeks of starvation, with three replicates totaling six samples, and so on. Sampling was conducted when the last group began to show spider death, and ultimately, the spiders died during the 19th week of starvation; thus, the last group was set as 19 weeks of starvation. In total, we obtained six groups of fat body samples and six groups of whole-body samples, totaling 36 samples.

### Transcriptome sequencing

A certain amount of RNA sample was taken and used to obtain mRNA from total RNA using oligo(dT). The mRNA was then fragmented, and random primers were subsequently used for cDNA synthesis. During the synthesis of the second strand of cDNA, dUTP was used instead of dTTP. The double-stranded cDNA was subjected to end repair, “A” addition, and adapter ligation. The enzyme UDG was used to digest the U-tagged second-strand template, followed by PCR and PCR product recovery. The library quality was assessed, and upon qualification, the product was circularized. The circular DNA molecules were subjected to rolling circle replication to form DNA nanoballs (DNBs) [55], which were then sequenced on the DNBSEQ platform.

### Genome assembly

Hifiasm v0.16.1 software was used with default parameters to perform an initial assembly of HiFi reads [23], resulting in contigs. The raw Hi-C reads were subsequently filtered using Hic-pro v3.1.0 [56], and the filtered Hi-C reads were subsequently analyzed with Juicer v1.6 to obtain a Hi-C interaction matrix [57]. 3d-dna v201008 was then employed to scaffold the contigs, yielding an initial pseudo-chromosomal genome [58]. Finally, manual corrections were applied using Juicerbox v1.11.08 to produce the final chromosomal genome.

### Repeat annotation

Repeat annotation consists of two parts, namely, utilizing an existing repeat library for repeat identification and constructing a repeat library from the genome itself for repeat identification, with the results from both parts being combined. RepeatMasker v4.1.2 [59] was used with the known repeat library Repbase v20181026 [60] for preliminary repeat identification. The construction of a self-derived repeat library was subsequently carried out using MITE Tracker v2.7.1 with default parameters to construct the mite library [61], followed by LTR analysis using ltrharvest v1.6.2 [62] and LTR_FINDER_parallel v1.1 [63], and the LTR library was integrated using LTR_retriever v2.9.5 [64]. RepeatModeler v2.0.2 was used for repeat analysis to obtain the repeat library [65]. The repeat libraries obtained from these tools were then integrated, and redundant sequences were removed using vsearch v2.23.0 to produce the final repeat library [66]. Repeat identification was then performed using the repeatmasker parameter (-lib) specifying this library, and the results from both identification processes were consolidated using the ProcessRepeats script included with RepeatMasker.

### Gene structure annotation

Initially, de novo gene structure prediction was performed using Augustus v3.4.0 [67] and SNAP [68]. The assembled transcripts were subsequently obtained using HISAT2 v2.2.1 [69] and StringTie v2.2.1 [70]. Transcriptomic evidence annotation was then conducted with PASA v2.5.2 [71]. Homology annotation was carried out using exonerate v2.4.0 [72] and GeMoMa v1.9 [73] with proteins from closely related species. Finally, the results from these three annotation methods were integrated using MAKER v3.01.04 [74] to produce the final annotation.

### GO functional annotation

Using BLASTP v2.12.0 (E value ≤ 1e-5) [75], spider protein sequences were aligned against homologous protein sequences in the UniProt Knowledgebase (UniProtKB) database [76], with the best alignments selected on the basis of bit score values. The GO annotations for the species were determined on the basis of the annotation information of the similar proteins. ID mapping was performed using the IDmapping file, where the first column represents the UniProtKB ID and the eighth column contains the GO annotations [77].

### KEGG annotation

KEGG annotation of genes was performed using KofamScan v1.3.0 [78], with the output format set to mapper and an e-value threshold of 1e-5. The required library configuration files included ko_list.gz (ftp://ftp.genome.jp/pub/db/kofam/ko_list.gz) and profiles.tar.gz (ftp://ftp.genome.jp/pub/db/kofam/profiles.tar.gz).

### Genome size and repeat correlation analysis

In addition to the *H. venatoria* genome, the genome sequences of a scorpion and 12 other spider species were collected and subjected to repeat analysis. The correlation between the proportions of different types of repeat sequences and genome size across the 14 species was calculated using the cor function from the R package stats v4.2.2 [79], with the analysis method set to Pearson. A correlation heatmap was generated using the corrplot v0.92 package [80].

### Phylogenetic tree construction

To reconstruct the evolutionary history of *H. venatoria*, a dataset comprising eight Araneae species and a Scorpiones outgroup (*Centruroides sculpturatus*) was utilized for maximum likelihood tree construction. First, we ran OrthoFinder v2.5.4 [81] to infer orthologs using BLASTP v2.12.0 [75] with a p value threshold of <1e-5, resulting in the identification of 1764 one-to-one orthologous sequences. Orthologs were aligned using MAFFT v7.520 [82] with the accurate option (L-INS-i) and trimmed using trimAl v1.4.rev15 [83] with the automated1 parameter. The trimmed alignments were then concatenated to serve as input for IQ-TREE v2.2.2.7 [84], using ModelFinder Plus (MFP) mode and 1000 bootstrap replicates.

### Gene family expansion and contraction analysis

We used CAFE5 v5.1.0 to investigate gene family expansion and contraction across selected species [85]. An ultrametric species tree was obtained using the MCMCTREE program in PAML v4.10.7 [86]. The calibration of the divergence time for species was derived from Magalhaes, Timetree (http://www.timetree.org/) and Paleobiodb (https://paleobiodb.org/) with Scorpiones stem (418-423 Mya), the split between *E. davidi* and seven other spiders (242-299 Mya) and the split between *U. diversus* and *H. venatoria* (173-240 Mya) [87, 88].

The gene family results were acquired from OrthoFinder, with only those families containing no more than 100 gene copies retained for further analysis. In CAFE5, the base model and unspecified Poisson distribution were used to conduct calculations. The significantly expanded families (p value < 0.05) in *H. venatoria* were analyzed for functional enrichment using the R package clusterProfiler v4.10.1 [89].

### Transcriptome analysis

Principal component analysis (PCA) was conducted using the R package FactoMineR v2.10 [90], and differential transcriptomic analysis was performed with the DESeq2 v1.42.1 package [91]. For the differential analysis of fat body stages (early, middle, and late), taking the early stage as an example, differentially expressed genes (DEGs) were identified for both the control (ck) and 2-week samples relative to the middle and late-stage samples. The logFoldChange threshold was set at 1, and a p value of 0.05 was considered to indicate statistical significance. The intersection of the two sets of DEGs was taken as the early-stage DEGs. The middle and late stages were analyzed similarly.

DEGs at the 19 W stage in the fat body and whole-body tissues were analyzed relative to other stages. Using the fat body as an example, DEGs were calculated for 19 W relative to the CK, 2 W, 4 W, 8 W, and 14 W periods. The intersection of these five sets of DEGs was considered to represent the DEGs at 19 W in the fat body. The logFoldChange threshold was set at 0.5, and a p value of 0.05 was considered significant for fat body differential analysis. For whole-body analysis, owing to the large differences in the transcriptome at 19 W compared to other periods, resulting in an excessive number of DEGs, the logFoldChange threshold was also set at 0.5, with a p value of 0.05 considered indicative of statistical significance.

WGCNA was performed on the fat body transcriptome via the R package WGCNA v1.72.5 [46], with the minimum module gene count set at 30 and the soft threshold power in fitIndices set at 10. For functional enrichment analysis, KEGG pathway enrichment was performed using the clusterProfiler v4.10.1 package, with a corrected padjust value less than 0.05 considered indicative of statistical significance.

### Cytochrome P450 family analysis

Three approaches were utilized to identify putative CYP genes. First, we downloaded known CYP amino acid sequences of five arthropods, namely, *Apis mellifera*, *Bombyx mori*, *Drosophila melanogaster*, *Pogonomyrmex barbatus*, and *Tetranychus urticae*, from the Cytochrome P450 Homepage (https://drnelson.uthsc.edu/). We used the protein sequences of eight Araneae species as queries to perform BLASTP v 2.12.0+ analysis against known CYPs, employing a threshold of p < 1e-30. Second, we used the hmmersearch program within HMMER v3.3.2 [92]. The P450 domain (PF00067) was searched against the candidate gene from the BLAST results, which required both full sequence and domain scores > 100. Finally, we manually removed the sequences that lacked any one of the five conserved motifs of CYPs (WxxxR, Gx[ED]T[TS], ExLR, PxxFxP[ED]RF, PFxxGxRxCx[GA] in Insecta).

For CYP family phylogeny, sequences of Araneae species and *Tetranychus urticae* were aligned with MAFFT (L-INS-i) and trimmed with trimAl (gappyout method). The trimmed sequences were then input to IQTREE2 (MFP mode), and a CYP phylogenetic tree was constructed. All trees presented in our manuscript were visualized using iTOL v6 (https://itol.embl.de/) [93].

### Synteny analysis

To assess the genomic synteny between *H. venatoria* and other species, we employed the JCVI v1.3.8 MCScan application for the calculations [94]. We have highlighted a specific clade within the CYP clan 3 members in the figure instead of a synteny block.

## Author Contributions

Guoqing Zhang, Yiru Wang and Hongcen Jiang prepared the sequencing samples and completed the genome assembly and gene annotation. Guoqing Zhang and Yiru Wang conducted the transcriptomic and evolutionary analyses. Guoqing Zhang wrote the initial draft of the manuscript, and Yi Wang supported the project and reviewed the manuscript.

## Acknowledgements

We thank Ling Xu at China Agricultural University, Sanyuan Ma at Southwest University for critical reading of the manuscript. National Key Research and Development Program of China [2023YFF0713900]; Science and Technology Innovation Key R&D Program of Chongqing (CSTB2022TIAD-STX0015), Special Fund for Youth Team and 2035 Pilot Plan for Innovative Research of Southwest University (SWU-XJLJ202306 and SWU-XDPY22009).

## Data availability

All high-throughput sequencing raw data, the genome assembly and gene annotation in this project were deposited into the Genome Sequence Archive (GSA) of the National Genomics Data Center (NGDC, https://ngdc.cncb.ac.cn/) and are available through BioProject ID PRJCA028207.

## Competing Interests

The authors declare no competing interests.

## List of Supplementary Information Provided

**Supplementary Figure S1. Length statistics of HiFi sequencing reads for *Heteropoda venatoria*, as well as the results of the genome survey.**

**Supplementary Figure S2. Transcriptomic analysis results of the fat body and whole-body in Heteropoda venatoria.** (A) Principal component analysis (PCA) results of the starvation resistance of all samples with whole-body. (B) Correlation analysis of the transcriptomes of the fat body and whole-body across various stages of starvation resistance. (C, D) The soft threshold of fat body and whole-body in WGCNA.

**Supplementary Figure S3. The results of WGCNA.** (A) The modules clustered of WGCNA in fat body transcriptomes. (B) The correlation between modules and traits. (C, D) Scatterplot of gene significance (y-axis) vs. module membership (x-axis) in blue and brown modules.

**Supplementary Figure S4. The expression and KEGG functional enrichment results of modules with WGCNA.** (A, B, C) The expression of blue, brown and turquoise modules. (D, E, F) The KEGG functional enrichment results of blue, brown and turquoise modules.

**Supplementary Figure S5. The analysis results of blue module in WGCNA.** (A) The expression trend of the four pathways enriched genes during the early, middle and late periods of starvation resistance. (B) The network of the transporter pathway-related genes enriched in blue module. (C) The network of four energy metabolism-related pathway (AMPK, TCA cycle, PPAR and insulin resistance) genes enriched in the blue module.

**Supplementary Figure S6. The expression trend of pathways enriched with DEGs in the fat body transcriptome of *Heteropoda venatoria* during a 19 W starvation period.**

**Supplementary Figure S7. Heatmap and KEGG pathway enrichment analysis of DEGs in the fat body transcriptome of *Heteropoda venatoria* after 19 weeks of starvation resistance.**

**Supplementary Figure S8. Phylogenetic tree of all P450 genes in eight spider species and *Tetranychus urticae*.**

**Supplementary Figure S9. The expression of P450 genes of fat body and whole-body across various stages of starvation resistance in *Heteropoda venatoria*.**

**Supplementary Figure S10. The expression of P450 genes of various tissues in *Trichonephila clavate*.**

**Supplementary Figure S11. The five common conserved motifs of all P450 genes and a specific sequence of CYP3 genes in *Heteropoda venatoria*.**

**Supplementary Figure S12. The seqLogo of conserved motifs of P450 genes in spiders.**

**Supplementary Table S1. The functional enrichment results of expanded family genes.**

**Supplementary Table S2. The functional enrichment results of blue, brown and turquoise modules.**

**Supplementary Table S3. The kegg annotation of Lysosome genes with blue, brown and turquoise modules.**

**Supplementary Table S4. The list of expanded family genes.**

**Supplementary Table S5. The DEGs of the fat body and whole-body transcriptomes at 19 W starvation period in *Heteropoda venatoria*.**

## Notes

### Competing Interest Statement

The authors have declared no competing interest.

