## Supplementary figures and images for "Genomic and transcriptomic analyses of *Heteropoda venatoria* reveal the expansion of P450 family for starvation resistance in spider"

### Supplementary Figure S1

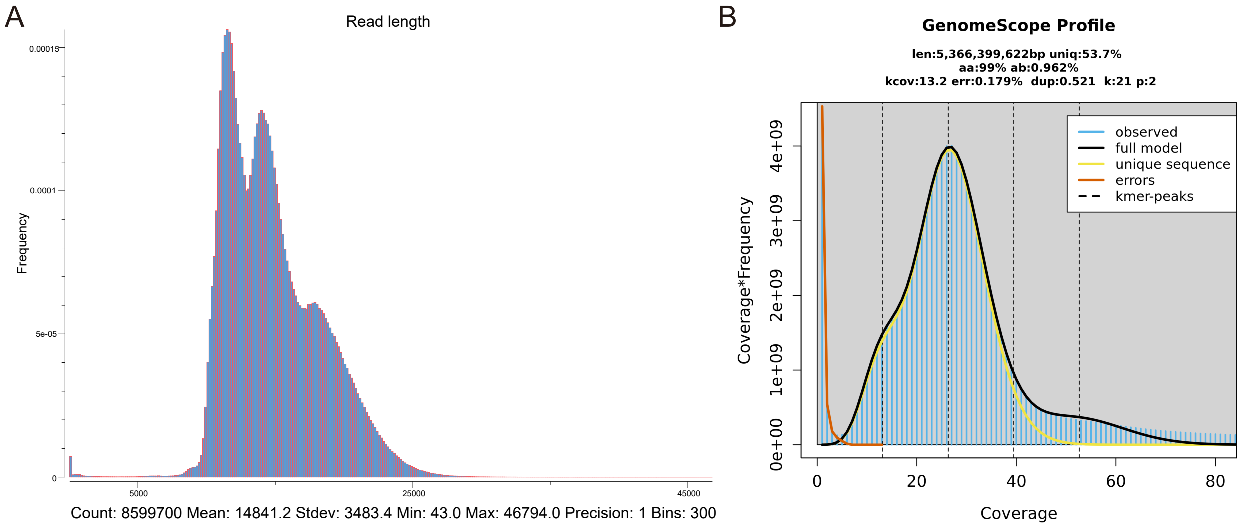

### Supplementary Figure S2

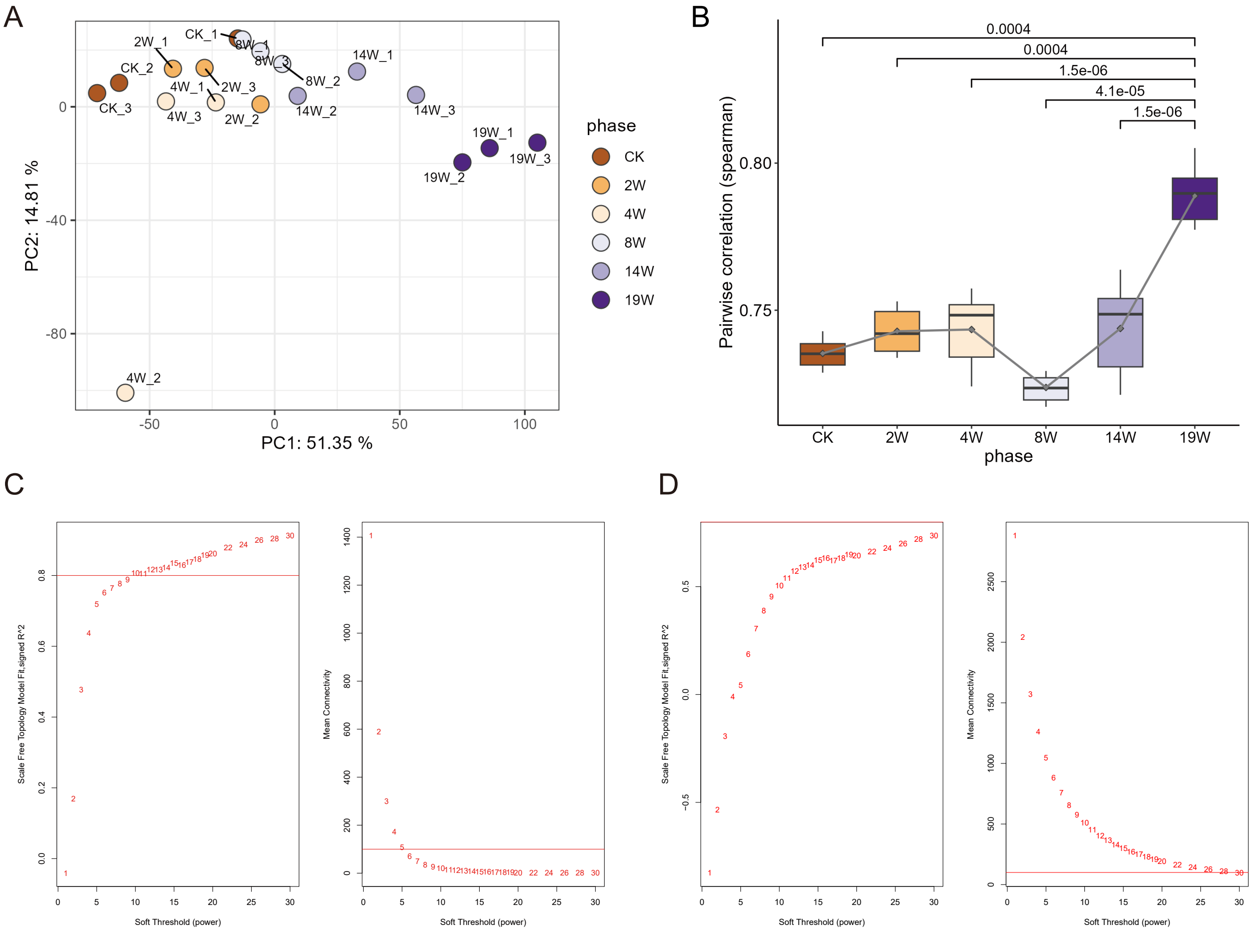

### Supplementary Figure S3

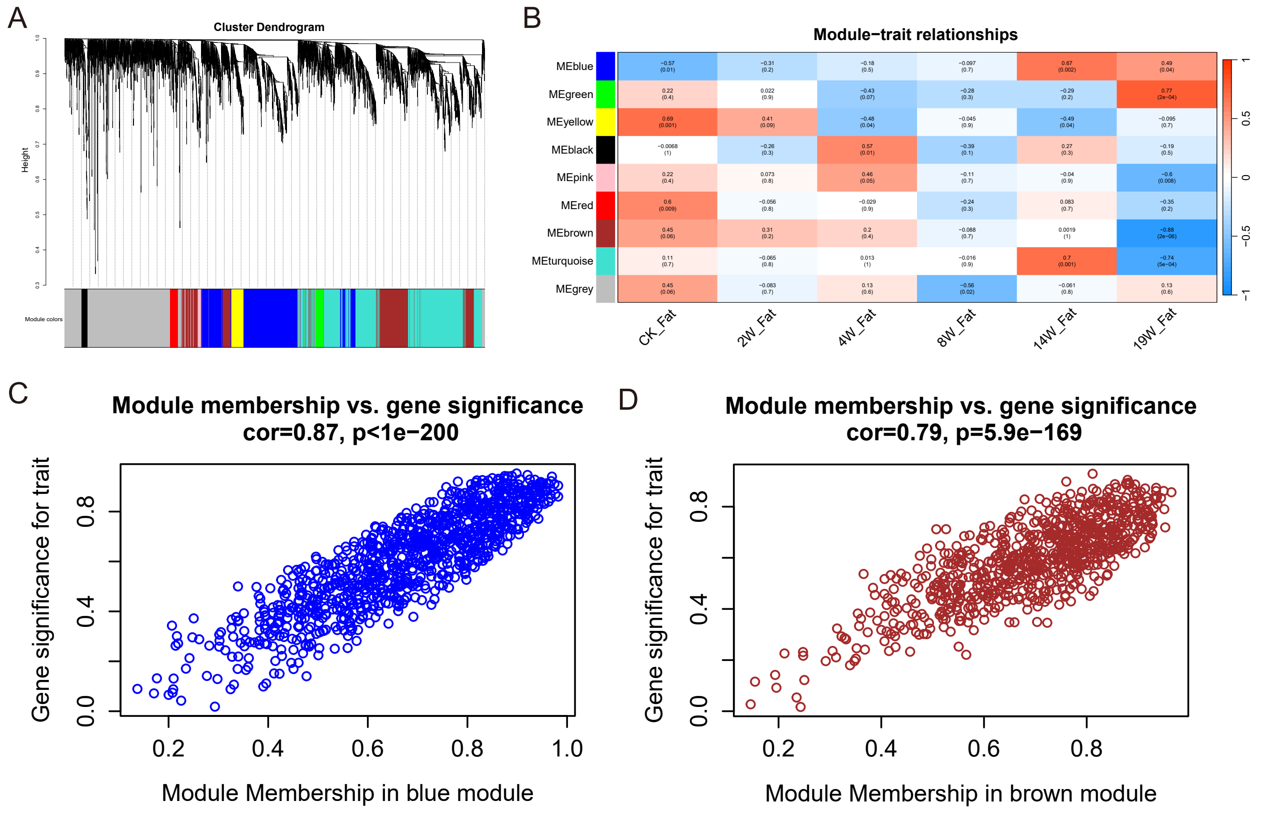

### Supplementary Figure S4

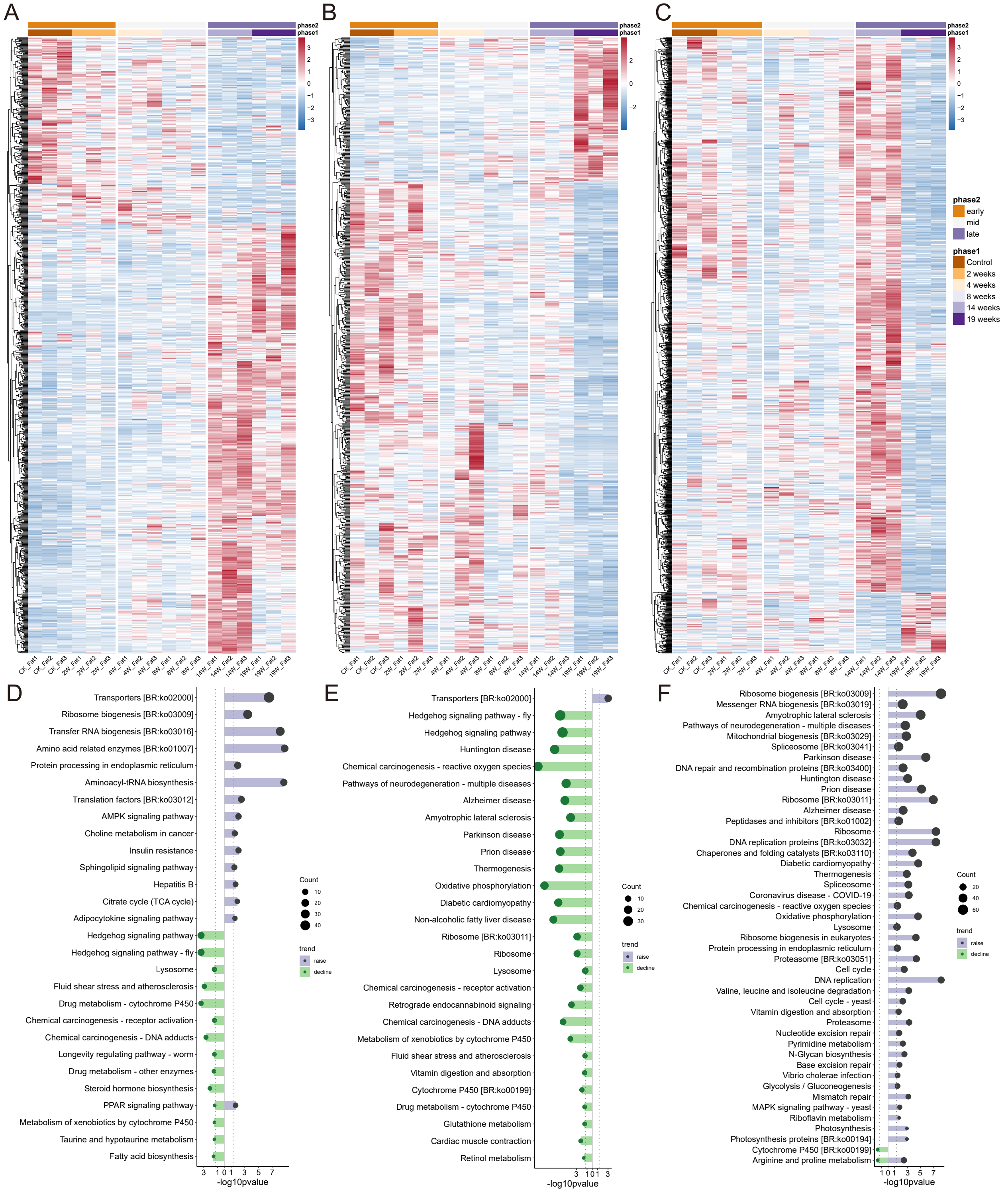

### Supplementary Figure S5

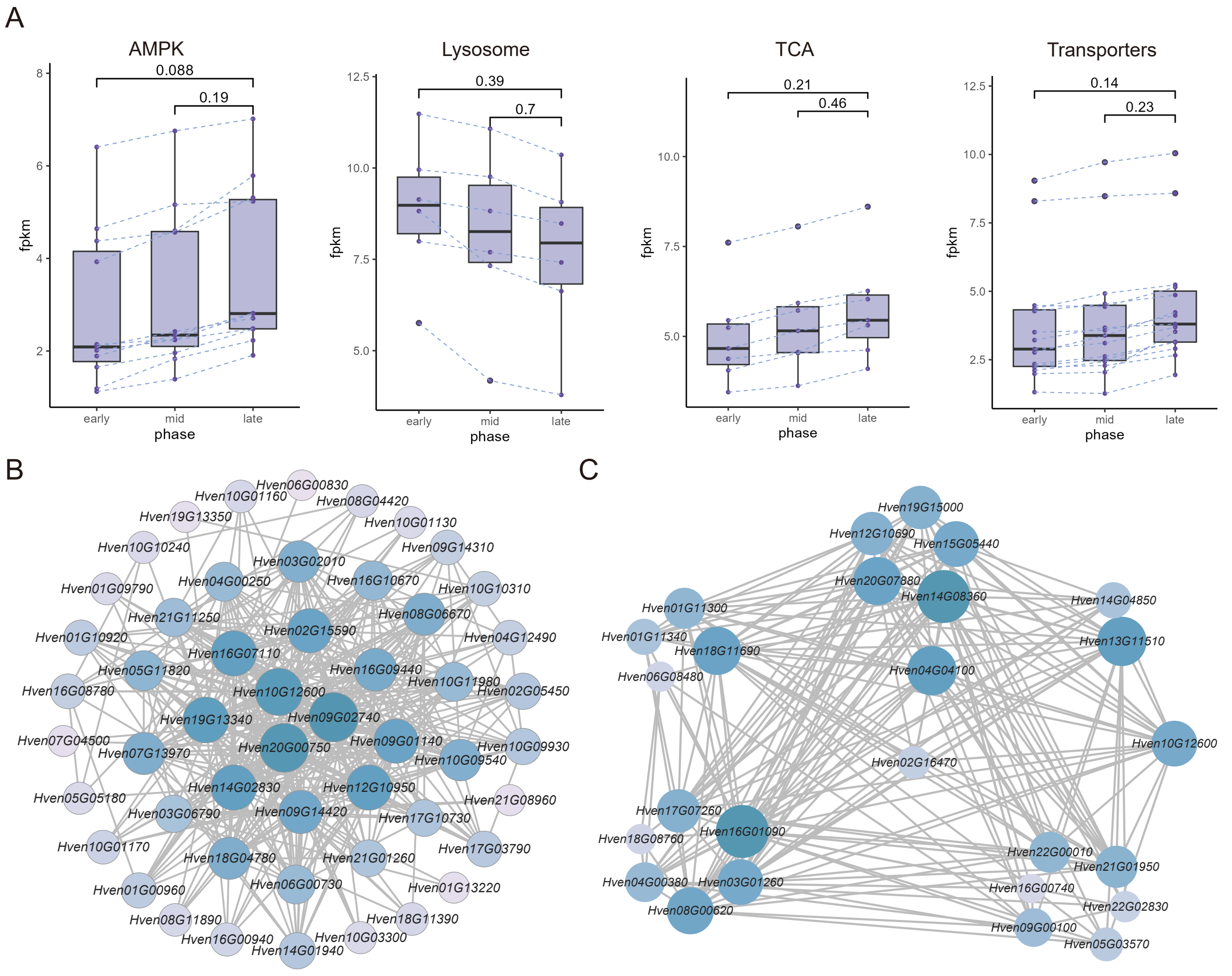

### Supplementary Figure S6

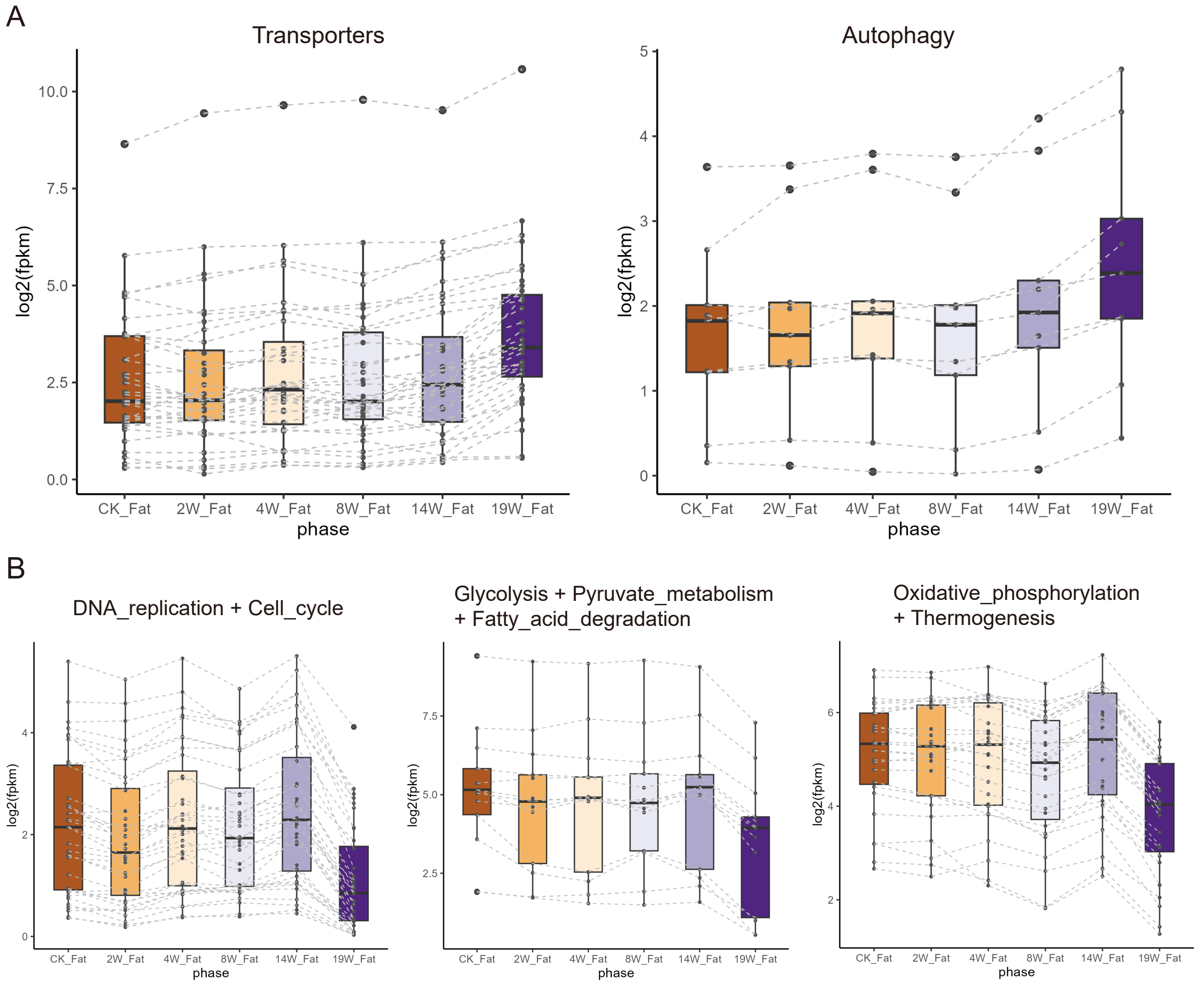

### Supplementary Figure S7

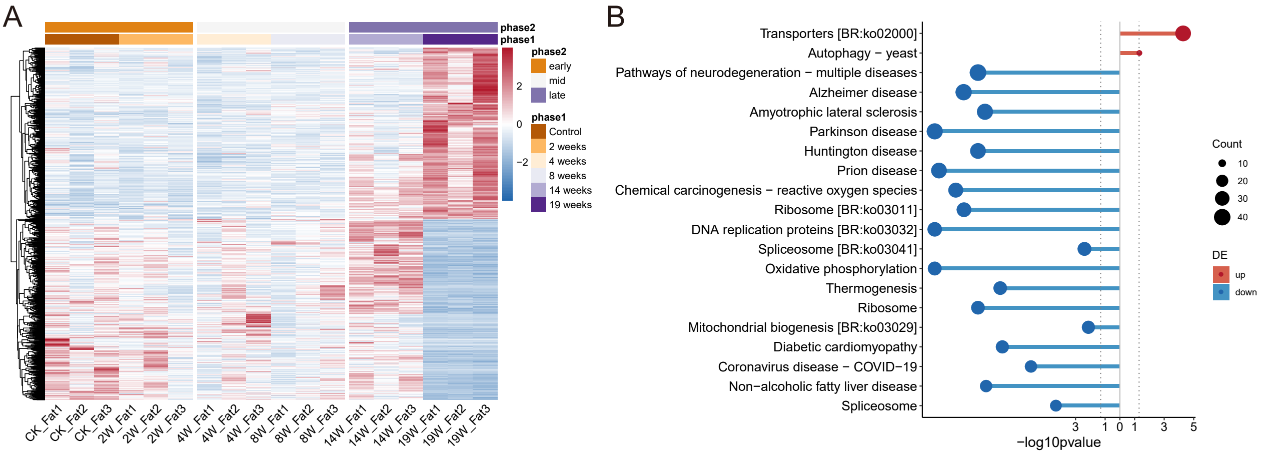

### Supplementary Figure S8

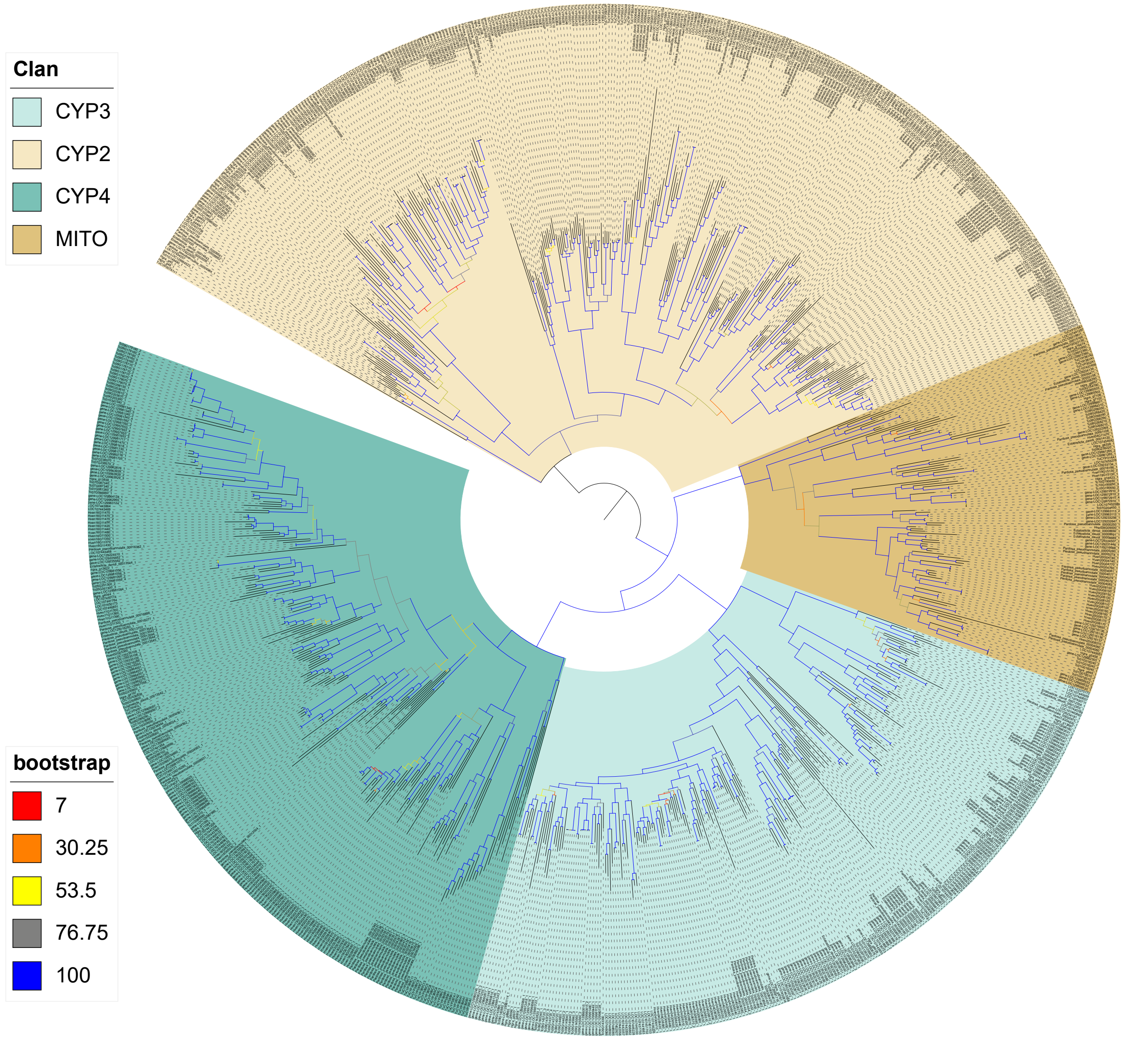

### Supplementary Figure S9

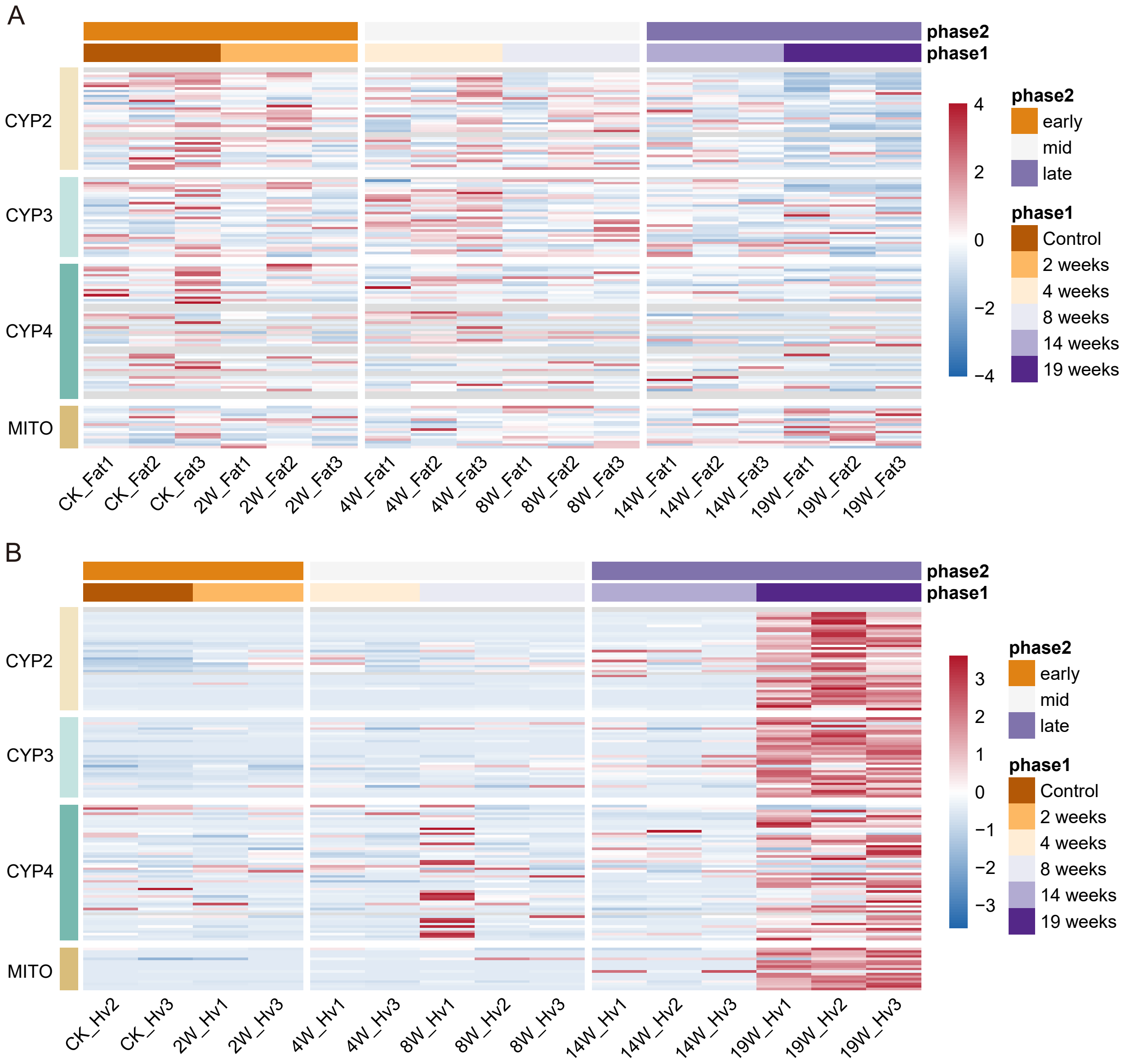

### Supplementary Figure S10

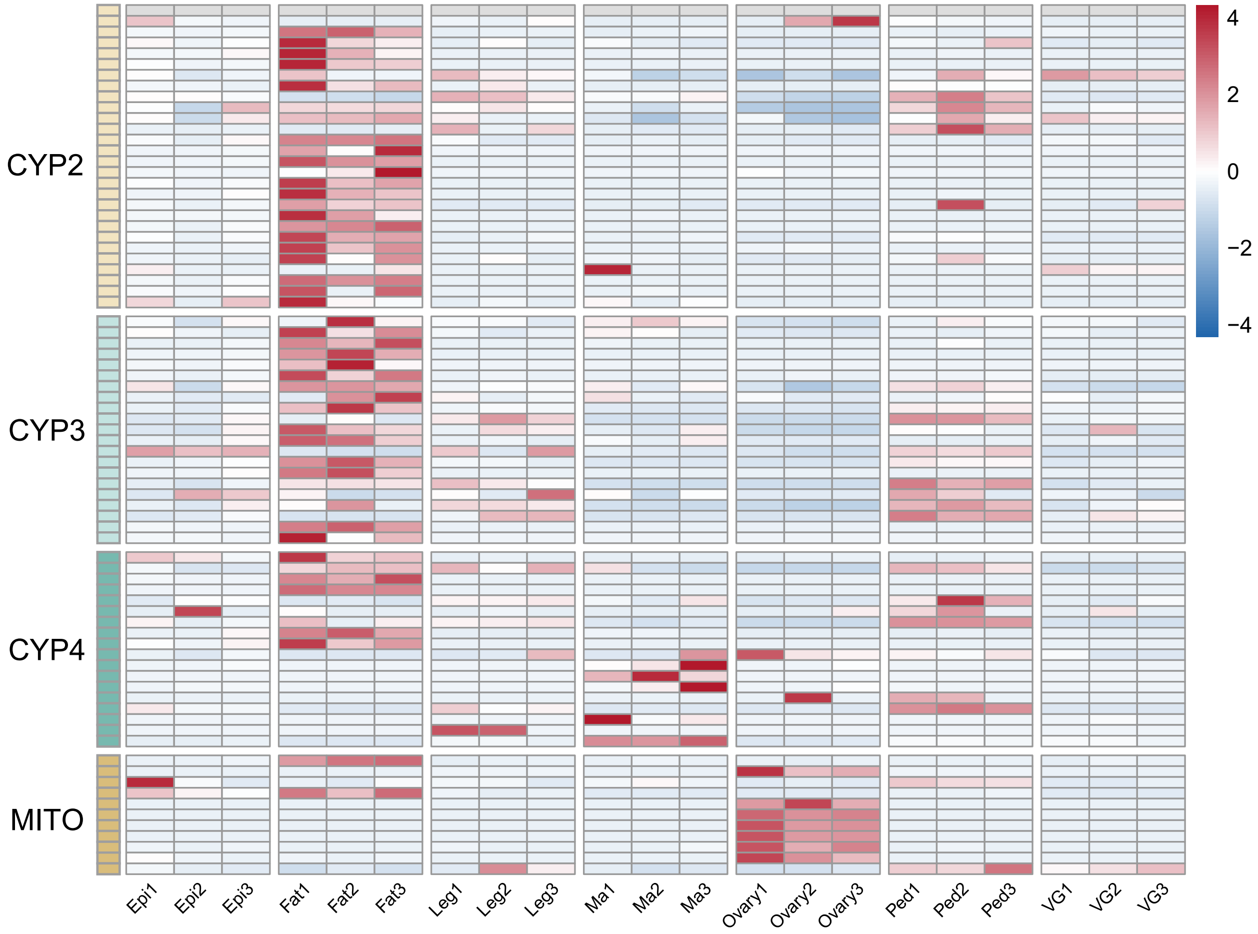

### Supplementary Figure S11

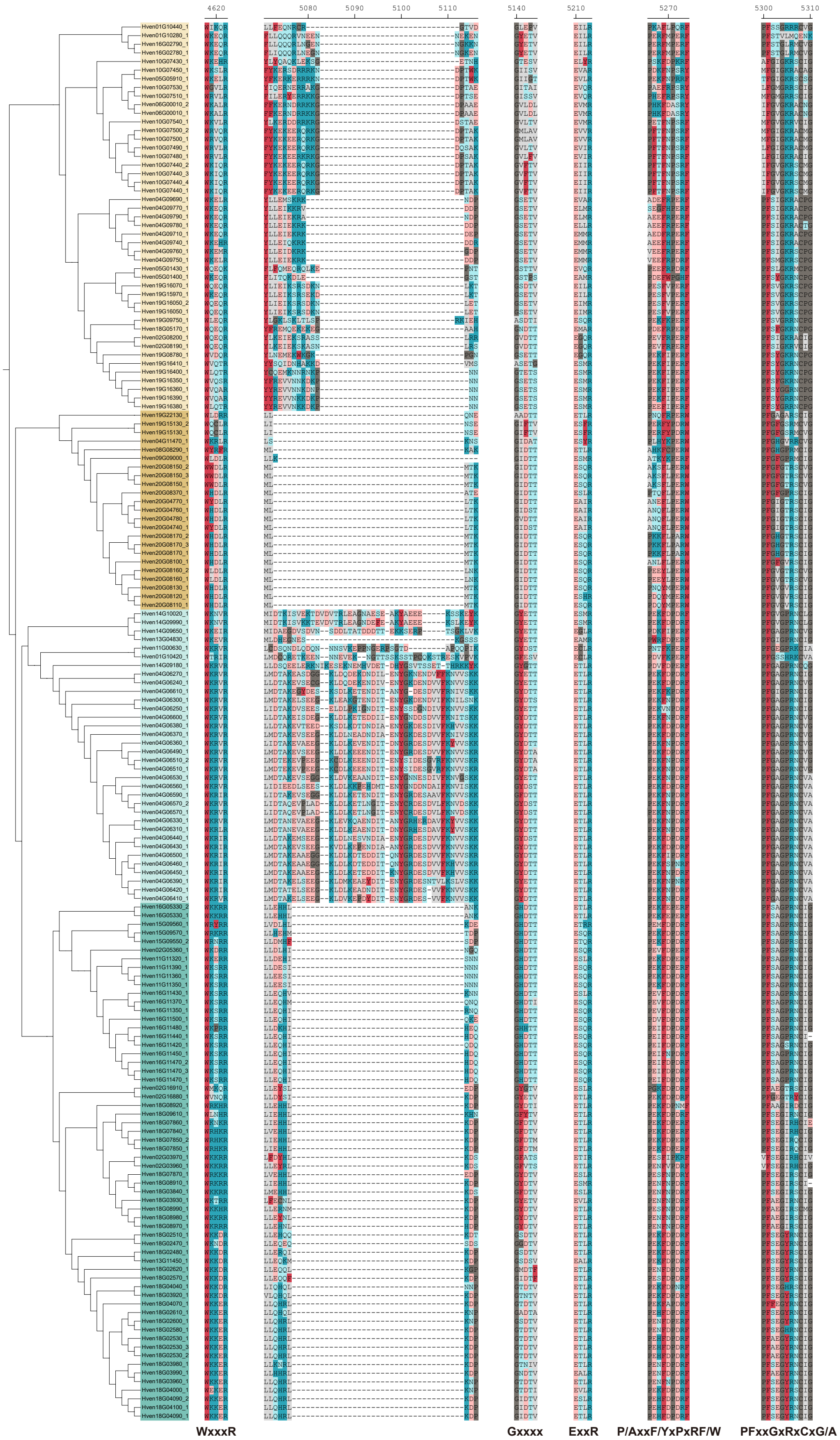

### Supplementary Figure S12

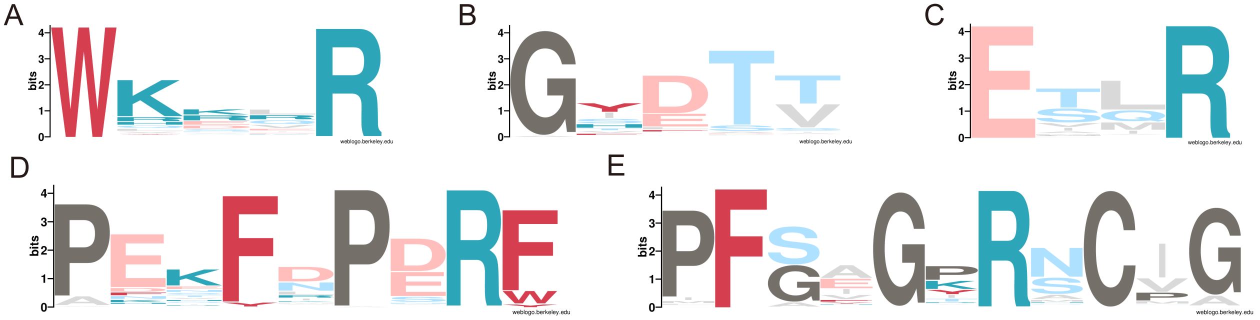
